# Full scale structural, mechanical and dynamical properties of HIV-1 liposomes

**DOI:** 10.1101/2021.03.24.436841

**Authors:** Alexander J. Bryer, Tyler Reddy, Edward Lyman, Juan R. Perilla

## Abstract

Enveloped viruses are enclosed by a lipid membrane inside of which are all of the components necessary for the virus life cycle; viral proteins, the viral genome and metabolites. Viral envelopes are lipid bilayers that adopt morphologies ranging from spheres to tubes. The envelope is derived from the host cell during viral replication. Thus, the composition of the bilayer depends on the complex constitution of lipids from the host-cell’s organelle(s) where assembly and/or budding of the viral particle occurs. Here, molecular dynamics (MD) simulations of authentic, asymmetric HIV-1 liposomes are used to derive a unique level of resolution of its full-scale structure, mechanics and dynamics. Analysis of the structural properties reveal the distribution of thicknesses of the bilayers over the entire liposome as well as its global fluctuations. Moreover, full-scale mechanical analyses are employed to derive the global bending rigidity of HIV-1 liposomes. Finally, dynamical properties of the lipid molecules reveal important relationships between their 3D diffusion, the location of lipid-rafts and the asymmetrical composition of the envelope. Overall, our simulations reveal complex relationships between the rich lipid composition of the HIV-1 liposome and its structural, mechanical and dynamical properties with critical consequences to different stages of HIV-1’s life cycle.

## Introduction

Enveloped viruses are molecular pathogens enclosed in a lipid bilayer acquired from the host cell during viral assembly and egress. ^1^ The lipid envelope and membrane proteins act as a container for all the components necessary to fulfill the virus life cycle, ^2^ adopting morphologies ranging from spheres to tubes. ^3^ The lipid bilayer itself is derived from the host cell during viral replication. Thus, the composition of the bilayer depends on the complex constitution of lipids from the host-cell’s organelle(s) where assembly and/or budding of the viral particle occurs.^4–7^ The human immunodeficiency viruses type 1 (HIV-1) is an enveloped retrovirus that infects immune system cells, specifically CD4^+^T cells and macrophages. ^8,9^ Initially, the virion enters the host cell by direct fusion with the plasma membrane, facilitated by viral membrane proteins. ^10,11^ During the late-stages of the life cycle, assembly of the virion inside the host-cell takes place at the plasma membrane directed by specific lipid-protein interactions. ^8,9,12^

Budding and assembly of the viral particle takes place at specific locations on the plasma membrane (PM).^5^ Essential to viral fitness is the structural Gag polyprotein which regulates several events during viral replication. ^9^ For instance, anchoring of HIV-1’s Gag to the inner leaflet drives viral particle assembly at specific microdomains of the PM. ^9,13^ More specifically, the anchoring of Gag to the PM is driven by the matrix domain (MA). MA contains a myristate moiety, that in the presence of phosphatidylinositol 4,5-bisphosphate (PIP_2_), facilitates the insertion of the acyl chain to the inner monolayer, inducing Gag assembly in PIP_2_-rich domains. ^14,15^ Moreover, the specific constitution of lipids plays a role in membrane deformation and curvature of the vesicle during particle budding,^16^ and the stiffness of the virion is an essential physical determinant for HIV-1 infectivity related to specific life-stages of the virus. ^17–19^

As previously metioned, HIV-1 virions are composed of proteins, metabolites, sugars and lipids. ^20^ High precision characterization of the lipidome of authentic HIV-1 virions has been reported by lipidomic analyses.^19,21,22^ In addition to sphingomyelin (SM) and cholesterol, the HIV-1 lipidome contains a high percentage of phosphatidylserine (PS) and phosphatidylethanolamine (PE).^18,19,21^ Therefore, it was initially suggested that the lipid composition of the viral lipid vesicle resembles that of a typical mammalian plasma membrane, with a prevalence of phosphatidylcholine (PC) and SM in the outer leaflet, and PS and PE lipids in the inner leaflet. ^23^ However, differences in the composition of the lipid vesicle and the plasma membrane have been observed, indicating that viral morphogenesis occurs on lipid microdomains of host-cells. ^7,9,22,24^

Although recent advances in experimental techniques allow the characterization of structural properties of entire viral particles, molecular dynamics (MD) simulations provide a unique level of resolution for full-scale virion models and large-scale macromolecular complexes. ^25–28^ Here, we present a full-scale model of the HIV-1 liposome with physiological lipid composition derived from lipidomics of infective HIV-1 virions. Structural and dynamical characterization of an authentic HIV-1 liposome were performed using massively parallel computing, ^29^ over 5.2 *μ*s of simulation. The model of the HIV-1 vesicle (Fig. 1) includes 24 distinct lipid-species (Fig. 2) with a composition asymmetrically distributed in the inner and outer leaflets (Tables S1 and S2).

**Figure 1:**
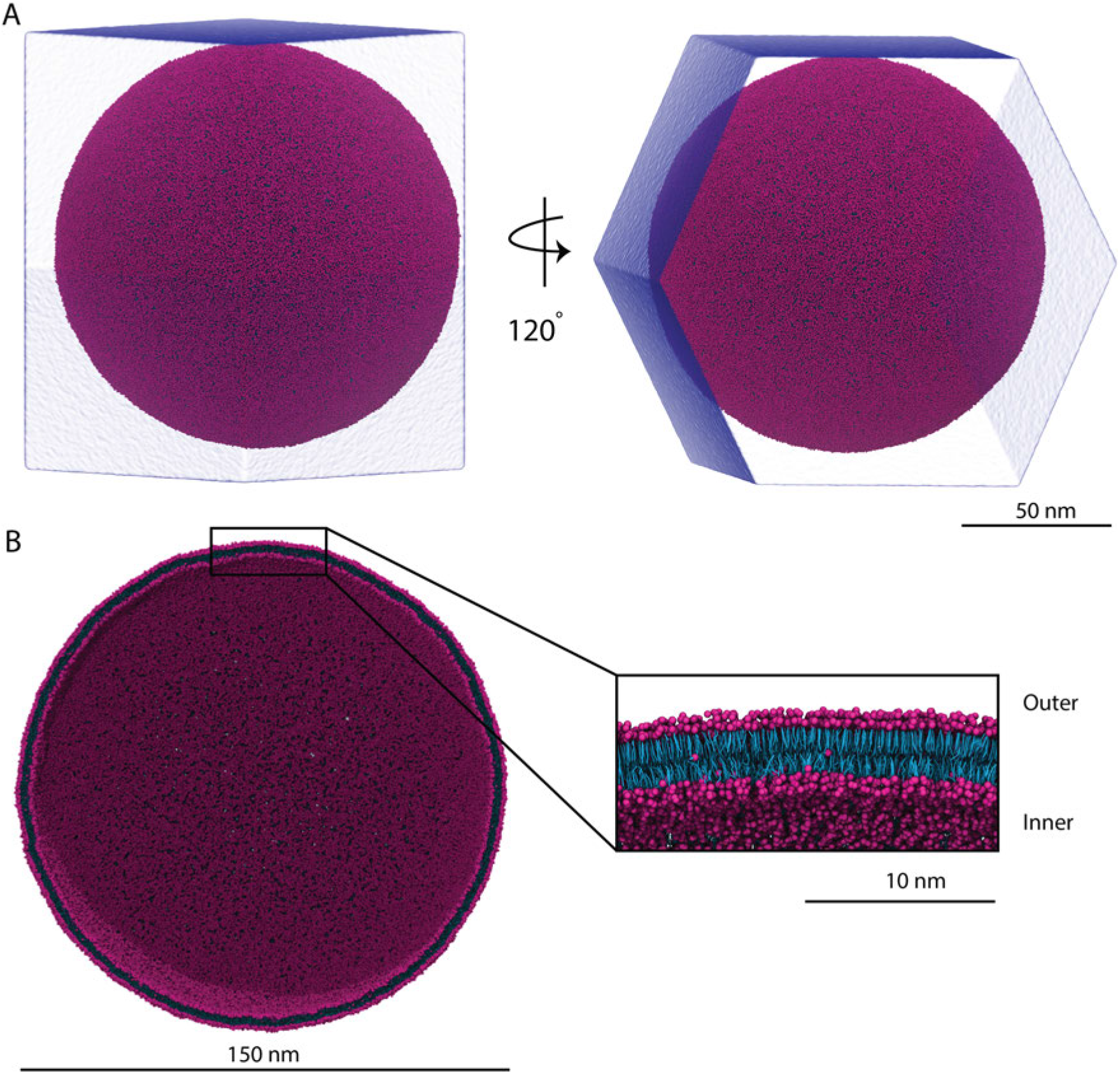
Full-scale model of a realistic HIV-1 lipid vesicle at united atom resolution (MARTINI force-field). **A** Full-view representation of the vesicle and solvent container used for simulation. **B** Clipped view of the vesicle. Each monolayer is shown in the inset. The headgroups are represented in magenta and tails are represented in cyan.

**Figure 2:**
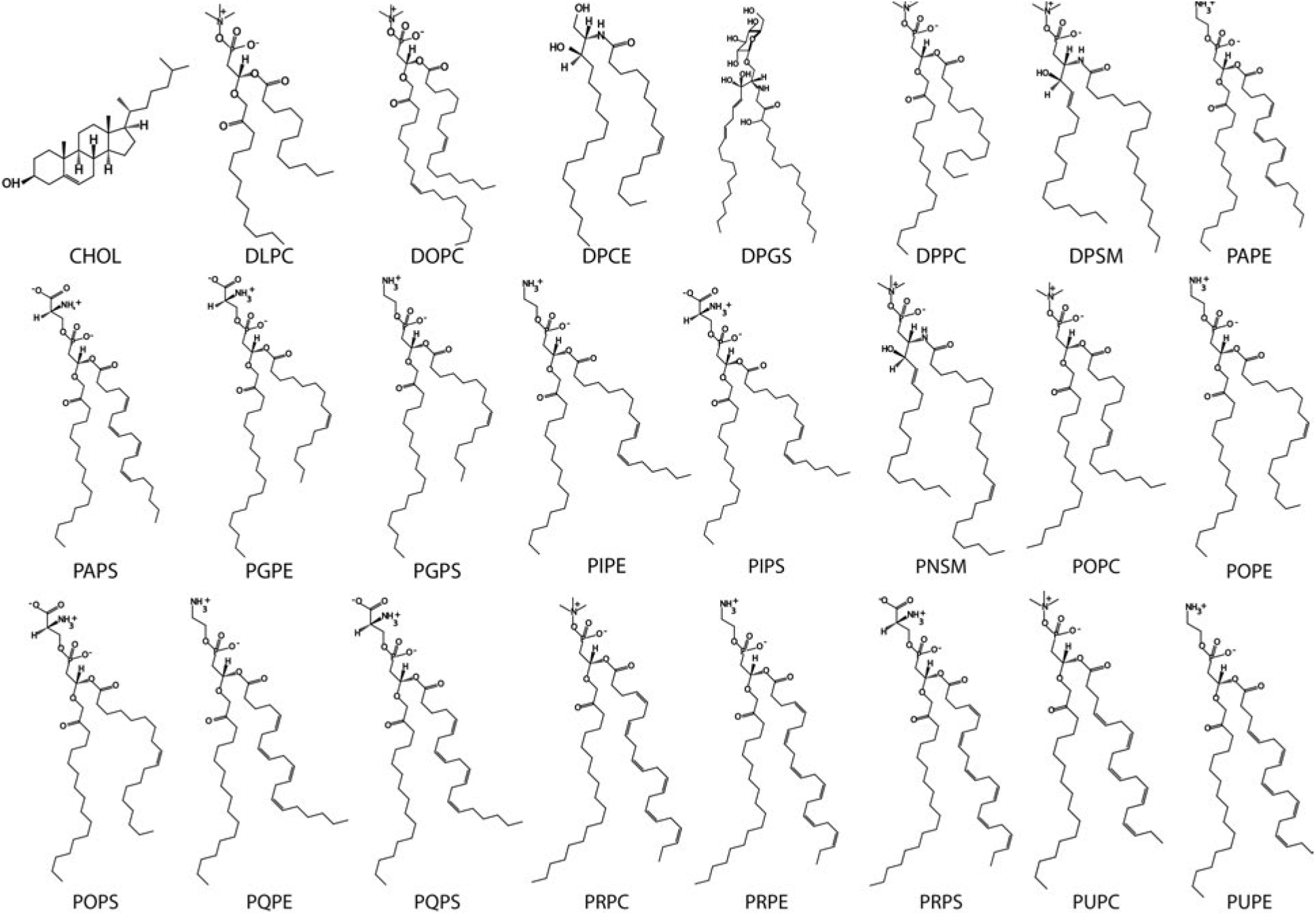
Chemical structures of the lipid species used in the composition of the HIV-1 lipid vesicle. A total of 24 lipid species were employed to match the complex composition found in authentic HIV-1 liposomes.

Despite significant flip-flop of lipids between the leaflets, the asymmetric composition is maintained within tight tolerances over the course of the simulation, suggesting a link between asymmetry and the shape of the envelope, even in the absence of envelope proteins. Analysis of the structural properties reveals the global fluctuations of the bilayer and of the bilayer thickness. The bending rigidity of the liposome is obtained from an analysis of fluctuations in its shape. We also find a heterogeneous organization of lipids into nanoscale ordered domains, with correspondingly heterogeneous dynamics. We anticipate that our results will guide the simulation of entire viral particles including membrane proteins, viral proteins and the viral genome; furthermore, the analysis framework derived herein can be readily applied to other enveloped viruses.

## Results

A full scale model of an HIV-1 liposome was built, equilibrated for 1.5 *μ*s and subsequently simulated for 5.2 *μ*s using the MARTINI force-field ^30^ (Movie M1). A rhombic dodecahedron solvation container was chosen to minimize the volume of the simulation system as shown in Fig. 1A. The liposome was constructed to have an initial outer diameter of 150 nm (Fig. 1B), falling in the range of outer lipid diameter values reported experimentally.^31^ The overall composition of the lipid vesicle was chosen to capture the chemical diversity of lipids observed by lipidomic analyses of infective HIV-1 virions (see Table S1).^19,21,22^ The initial distribution of lipids across the two leaflets is not known precisely for the HIV-1 virion, thus it was chosen based on well known plasma membrane asymmetry: negative charge, unsaturation, and ethanolamine on the inner leaflet; sphingolipids on the outer leaflet. ^24^ From the simulations, a wealth of biophysical properties were determined for an authentic viral liposome as presented in the following sections.

### Bilayer thickness and liposome sphericity

The outer diameter and sphericity of the HIV-1 lipid vesicle was measured over the first 1.5 *μ*s of equilibration. Convergence is observed for both parameters, with an average of 145 nm and 0.9975 for the outer diameter and the sphericity, respectively (Figure S1). During production, an increase of the outer diameter from 145 nm to 147 nm is observed. Sphericity is maintained with an average of 0.9984 throughout 5.2 *μ*s production sampling (Figure S2).

Estimation of the bilayer thickness was obtained from uniformly distributed regions of the vesicle; uniform sampling was achieved using a Poisson disk sampling technique ^32^ (blue points on Figure 4 **A**). Uniformly distributed disks of 1 nm in diameter were generated, small enough to minimize the effect of membrane curvature in the estimation of centroids on each of the leaflets. The bilayer thickness was initially calculated for the last frame of the vesicle simulation and the observed thickness values are shown to range from 34 Å to 36 Å, measured from headgroup to headgroup, with only small fluctuations, on the order of 1 Å of the bilayer thickness, are observed for the entire vesicle (Figure 4**B**,**C**).

**Figure 4:**
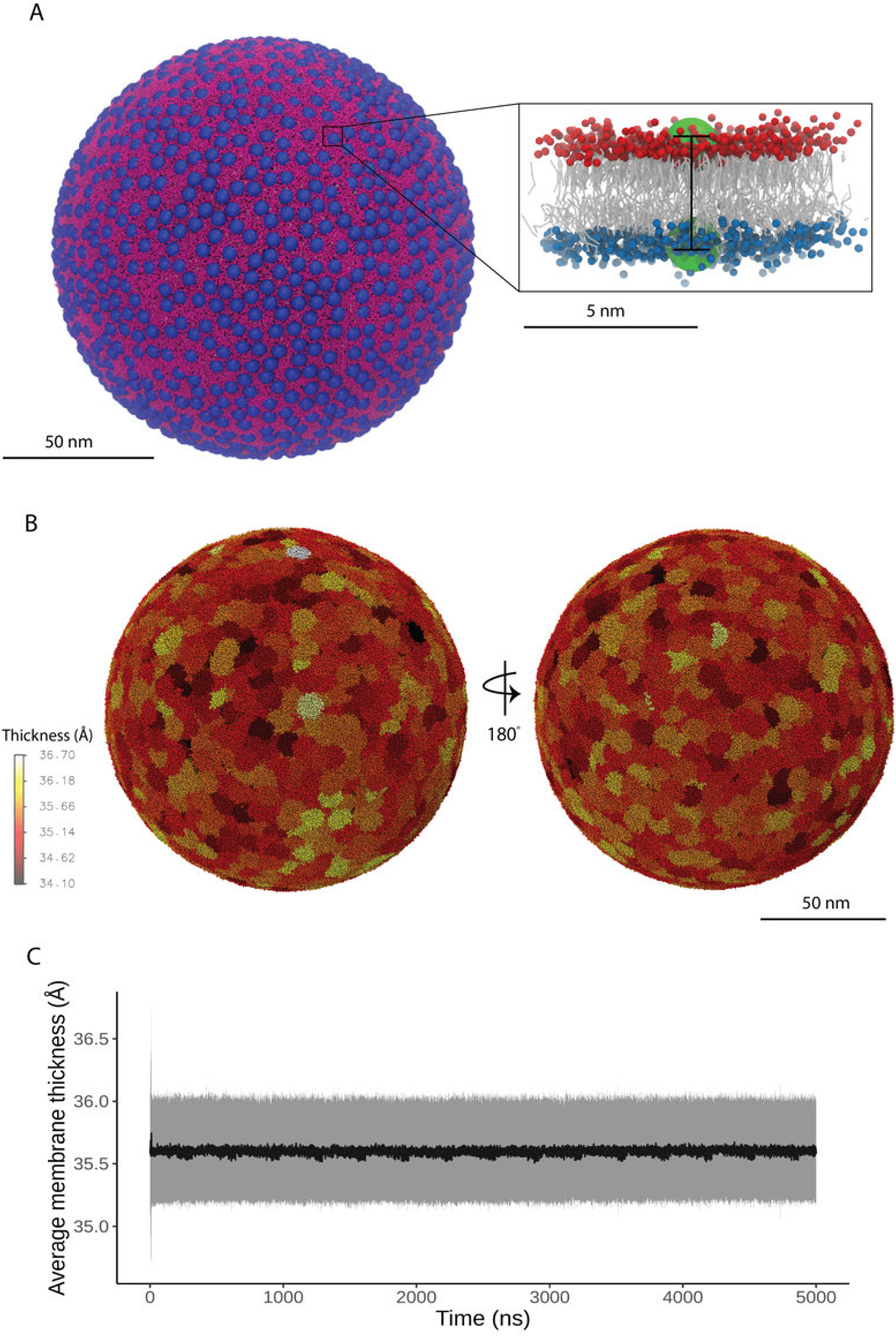
Local bilayer thickness of the HIV-1 lipid vesicle. **A** Uniform sampling representation over the surface of the vesicle (dark blue spheres). In the inset, lipids in the outer and inner monolayer are shown in red and blue, respectively, and the centroids of the particle selection are shown as green spheres for visual clarity. The distance between these two coordinates is used to deduce the local bilayer thickness. **B** Representation of local membrane thickness values over the surface of the vesicle. The visualization corresponds to the last frame of MD production. **C** Average bilayer thickness over the course of 5 μs of MD production. The standard deviation of each measurement is represented in silver.

Subsequently, thickness analyses were extended to the entire simulation (using VMD MPI and massive parallel computing to enhance performance ^29,32^) yielding an average bilayer thickness of 35.5 ± 0.4 Å with normally distributed variations in thickness (Figure S5). The observed thickness and its variation is in agreement with the 35 Å thickness observed by cryo-electron tomography for vesicles with cholesterol molar ratio of 42%. ^33^ In order to see whether thickness variations are caused by variations in composition, the compositions of three different thickness ranges at *σ*, 2*σ* and 3*σ* were determined. No relationship between the local membrane thickness and its composition was found. (Figure S5).

### Transmembrane asymmetry

Significant diffusion of lipids between the bilayer leaflets (“transverse” diffusion) is observed throughout the simulation (Figure 5, S3; Movie M2). To characterize the transverse diffusion of lipids, flip (outer-to-inner) and flop (inner-to-outer) rates were calculated separately during the 5.2 *μ*s of MD production for each lipid species. Previous MD simulation on flat-patch asymmetric membranes suggest the flip and flop rates are slow processes in the second timescale, supported by experimental small-angle neutron scattering (SANS) results. ^34^ The rate of flip and flop are influenced by the chain length, temperature and chemical character of the lipid headgroup.

**Figure 5:**
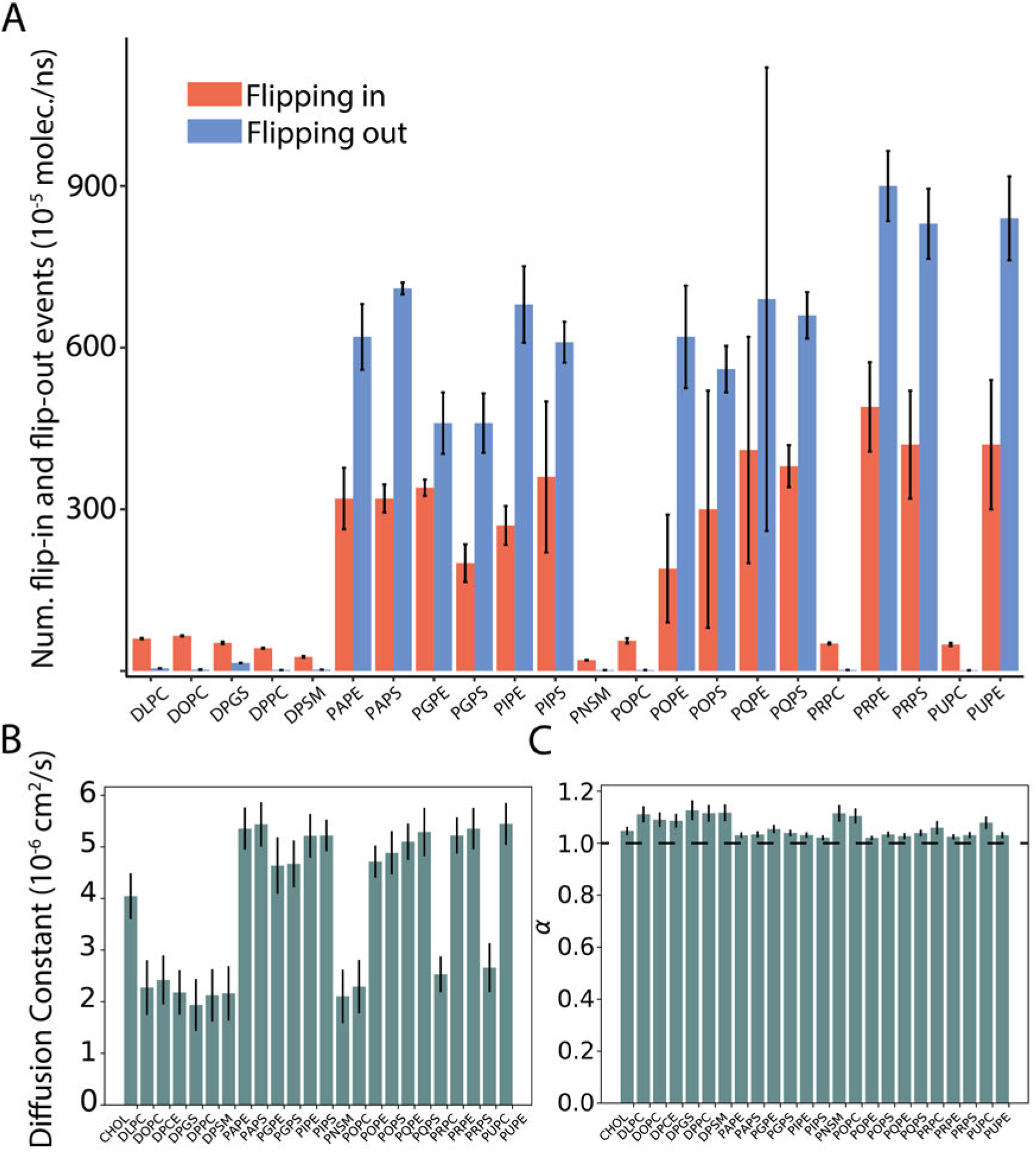
Diffusion of lipids on the complex vesicle model. **A** Trans-bilayer rates of lipids present in the vesicle after 5.2 *μ*s of MD production. The average outer to inner rate (blue) and outer to inner (red) rates were estimated by calculating the cumulative number of flipping events per lipid species from three different starting points during the simulation: 0, 200 and 4,000 ns. The values reported are normalized per number of events per molecule. **B** Lipid lateral diffusion coefficient and **C** scaling factor. The average is reported at windows sizes of 25, 50, 100, 250, 500 and 1000 ns

Average flip-flop rates were calculated for each lipid species over the entire production simulation of the HIV-1 liposome (Figure 5**A**); transbilayer flip-flop rates over different portions of the simulation are shown in Supplemental Figure S3). There are clearly two classes of lipids in Figure 5 **A**: lipids with rates less than 100(10^−5^) events/ns*molecule, and lipids with rates higher than 200(10^−5^) events*/*ns * molecule. The first category corresponds exclusively to lipid types that are enriched in the outer leaflet (sphingolipids and saturated chain lipids), while the second is exclusively lipids enriched on the inner leaflet (highly unsaturated lipids with PE and PS headgroups). Although the flip-flop rates are slow (as expected for lipid transbilayer rates), they do represent a substantial lipid translocation in the aggregate for many lipid types. For example, the total number of PAPS inner→ outer events is 40, while the total number of outer → inner events is 15. In spite of this, the initial and final numbers of PAPS in each leaflet are extremely asymmetric, and maintained within ± 2 lipids. Indeed, the total number of flipping events is larger than the number of PAPS in the outer leaflet. This is true for many of the lipid types, and indicates (i) that the difference in flip flop rates observed between inner and outer leaflets is not due to directional transport as the composition equilibrates by flip flop, and (ii) that compositional asymmetry would be maintained over longer timescales. Given that the composition of the two leaflets has not been experimentally determined, it is remarkable that the initial asymmetry is maintained during the *t* = 5.2 *μ*s of MD production, despite significant rates of lipid flip flop.

Given the observed stability of lipidomic asymmetry, the leaflet compositions reported in Figure 3 yield a good model for an HIV-1 lipid vesicle. These results are not only in agreement with bulk lipidomics results for the HIV-1 virion, ^21^ but are also supported by lipidomics experiments that suggest the formation of lipid domains on the host-cell, where budding and release of the viral particle takes place. ^35,36^

**Figure 3:**
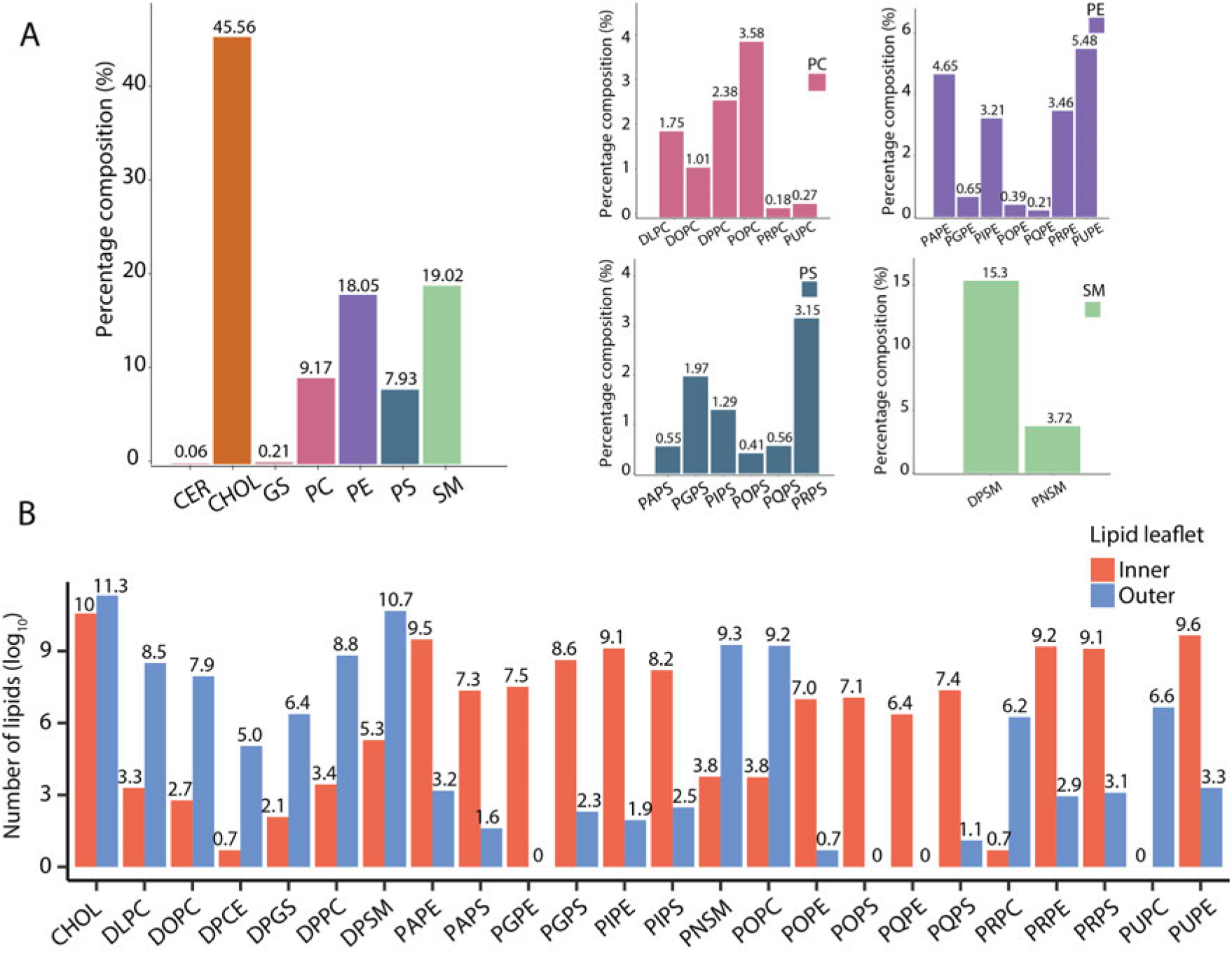
Lipid composition of an authentic HIV-1 liposome. The chemical composition is represented by lipid species breakdown and monolayer breakdown. The lipidome of the model consists of ceramides (CER), cholesterol (CHOL), glucosylceramides (GS), phosphatidyl-choline (PC), phosphatidylserine (PS), phosphatidylethanolamine (PE) and sphingomyelin (SM). **A** Breakdown percentage composition by lipid species. On the right side, the break-down is shown by lipid type; and on the left side, the lipid type breakdown composition is exhibited for lipid types with more than one species present in the vesicle. **B** Lipid composition of each monolayer after 5.2 μs of MD production.

### Lateral diffusion of lipids is heterogeneous within and between the leaflets

Characterization of the lateral diffusion of lipids was performed using methods previously developed by the authors for the analysis of full-scale virions.^27^ Analysis of time- and ensemble-averaged lipid mean squared displacements (MSD) show the diffusion to be Fickian (not subdiffusive). The diffusion constant (*D*) for different lipids in the inner and outer leaflets was obtained from fits of the MSD (Figure 5 **B**), with all diffusion constants on the order of 10^−6^ cm^2^/s.

Upon visual inspection of the trajectories it is clear that there are lipid domains on the vesicle that displace laterally much more slowly than the surrounding lipid matrix (Movie M3). To characterize these lipid regions based on their lateral mobility, the root-mean-squared fluctuations were calculated for each lipid molecule over different windows of the final 500 nanoseconds of the production simulation (Figure 6). In figure 6 **A**, several localized domains of low mobility lipids (dark patches) are observed, with diameters of roughly 10 nm. No such domains are observed on the inner leaflet (6 **B**). A further characterization of some of the most dramatic low-mobility domains on the outer monolayer reveals that these domains are enriched in cholesterol, sphingomyelin and phosphocholine (Figure 6 **B C**).

**Figure 6:**
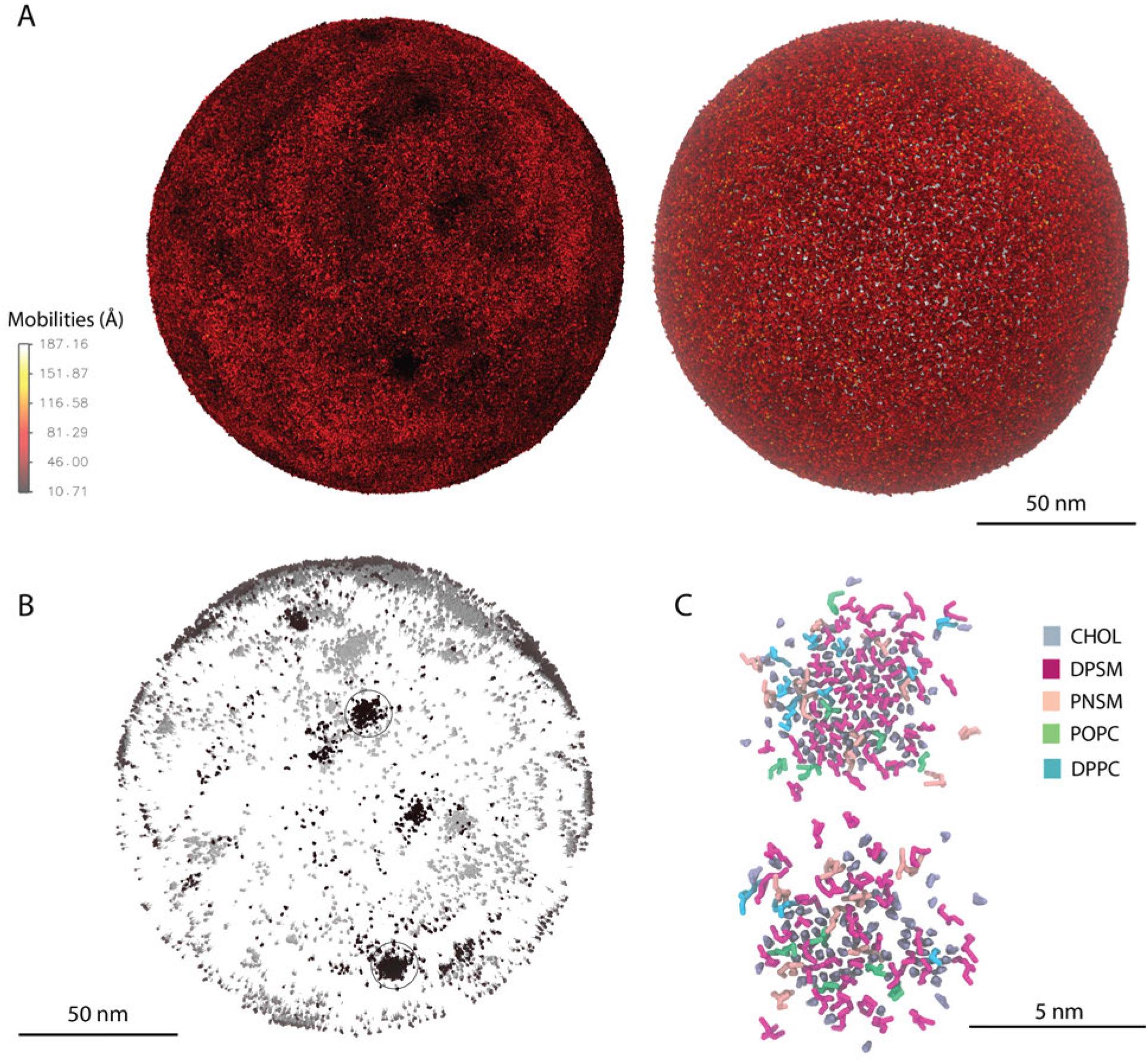
Mobility analysis of lipids and lipid regions. **A** Mobility of lipid headgroups after 1 μs of production MD. Headgroups in the outer leaflet (left) and the inner (right) leaflet show distinctive mobilities. **B** Determination of low mobility lipid regions (group of lipids with mobilities lower than 20 Å in the outer leaflet of the vesicle). Domains of aggregated lipids are characterized and classified (black circle). **C** Lipid composition of low-mobility lipid-regions in the outer leaflet. Lipids are colored according the lipid type.

The low mobility domains remain intact for a few nanoseconds and subsequently diffuse over the surface of the vesicle – they are transient, fluctuating domains which appear to be intrinsic to the composition of the outer leaflet of the lipid vesicle. During the simulation these low-mobility regions appear and disappear, highlighting their dynamic nature (Movie M3). Importantly, these low mobility regions are only observed in the outer leaflet and completely absent in the inner leaflet S7.

### Elastic properties

The bending rigidity *k*_*c*_ ^37,38^ of a flat bilayer can be obtained from spectral analysis of the fluctuations of the height of the membrane above a reference plane. However, the HIV-1 lipid vesicle is a quasi-spherical membrane that remains intact during the simulation, and therefore the instantaneous shape of the vesicle is decomposed into spherical harmonics ^38^ (Fig. 7).

**Figure 7:**
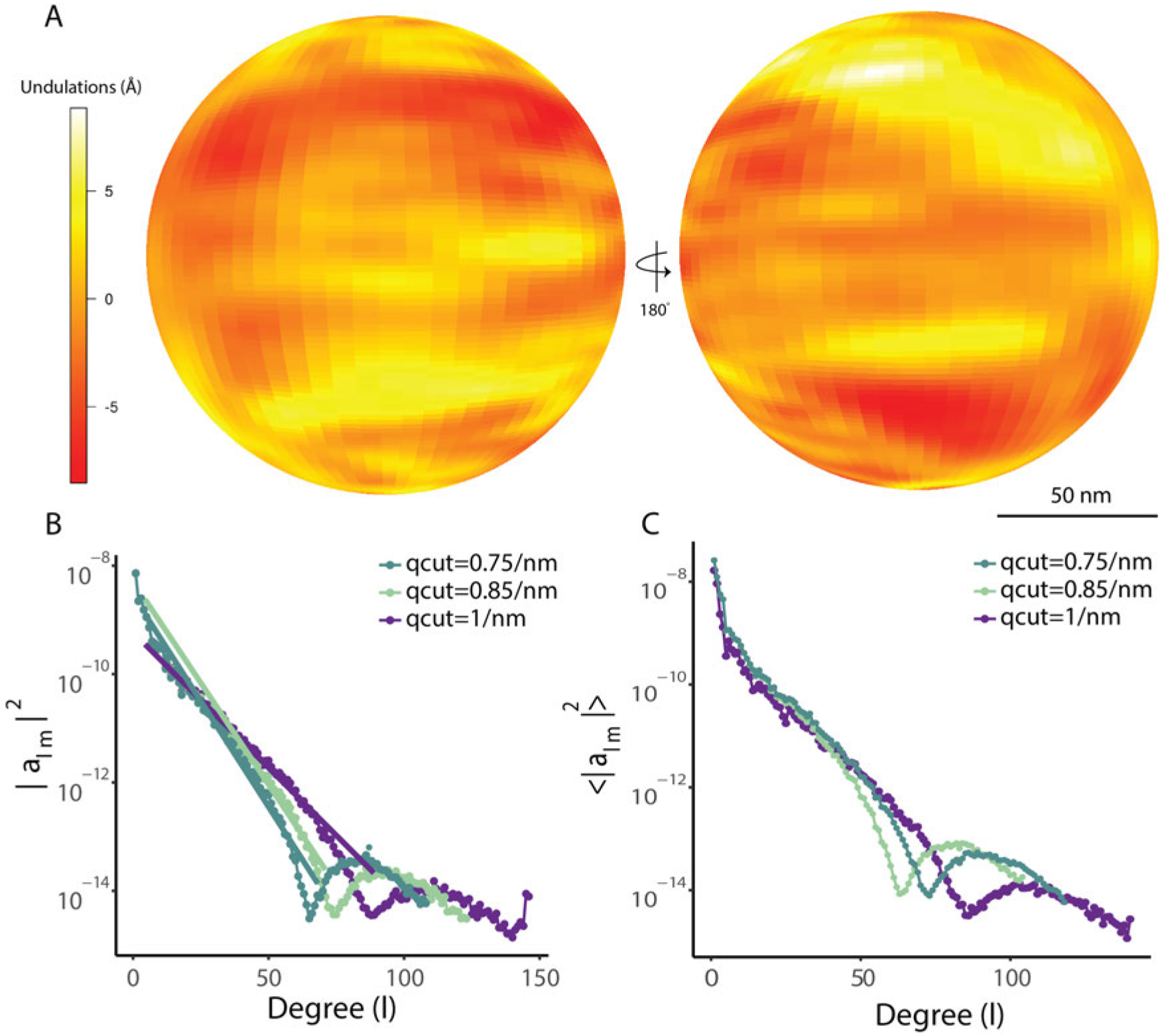
Bending rigidity of the HIV-1 lipid vesicle. **A** Undulating surface of the radial fluctuations as obseverd after 5 μs of MD production. **B** Undulation spectra from the expansion in spherical harmonic of the undulating surface for a single simulation snapshot of the liposome; the fit is shown for wave-number *q* cutoffs^38^ of 0.75 nm^−1^, 0.85 nm^−1^ and 1 nm^−1^. A linear fit is shown for the initial l-degrees for the three different cutoffs. **C** Ensemble average undulation spectra of the HIV-1 liposome. The bending rigidity is estimated from the Helfrich continuum model ^38^ for bilayer undulations yielding a value of 109*k*_*b*_*T* at 298K.

Following Braun and Sachs, ^38^ the bending rigidity of the viral liposome was obtained from the power spectrum of shape fluctuations in the spherical harmonic basis. The coefficients for the spherical harmonic expansion were obtained from a nonlinear fit of degree 74 and order 149. The undulation power spectrum in Figure 7 **B** was then computed by fitting *a*_*lm*_ to the power spectrum model in equation 5, yielding *k*_*c*_ = 109 *k*_*b*_*T* at 298K. The bending rigidity obtained in the present study implies that the HIV lipid vesicle remains flexible during simulation.

Flexible envelopes are characteristic of mature HIV-1 virions. Flexibility facilitates entry of mature virions, whereas immature virions exhibit a stiffer viral envelope which cannot enter host cells efficiently.^17^ Despite the presence of the low-mobility ordered domains described above, the viral envelope is still clearly in a fluid phase, as the nanoscale ordered domains are transient, and the bending rigidity is still significantly below values reported in the literature for a membrane below the main phase transition temperature. ^39^

The composition of the outer leaflet of the HIV-1 liposome (rich in sphingolipids and cholesterol) is consistent with raft-like regions of the plasma membrane, from which HIV-1 is believed to escape the cell. In simpler model membrane mixtures, the liquid-ordered phase has several features in common with rafts–it is enriched in high melting temperature lipids (like SM) and cholesterol, the hydrocarbon chains are ordered, and therefore the bending modulus is likely higher than in single component fluid membranes. ^40,41^ Recently, Weiner, et al. demonstrated liquid-ordered domain formation in all-atom simulations of more complex mixtures, designed to mimic the asymmetry of the plasma membrane. ^42^

The transient ordered regions observed in the outer leaflet of the HIV-1 liposome shown in Fig. 6 are also observed in bilayer simulations of the liquid-ordered phase. ^43^ Taken together, the analysis of the HIV-1 liposome simulation indicates that the outer leaflet of the liposome is in an ordered or raft-like phase. The fact that the composition is heterogeneous across the surface of the vesicle suggests that viral assembly might exploit fluctuations in lipid composition to organize surface proteins for critical functions like viral fusion and entry, as suggested previously for influenza. ^44^ Indeed, Tamm and coworkers have shown that gp41 mediated fusion occurs in heterogeneous membranes at the edge of cholesterol-rich domains. ^45^

## Discussion

We have performed a full-scale simulation of a complex, realistic vesicle at the united atom particle-based level. The united atom system was built from experimental data, introducing lipid asymmetry in the compositions of the outer and inner leaflet. The results from equilibration show the vesicle maintains sphericity reported by experimental techniques. In addition, the lipid asymmetry is maintained during the MD production suggesting our model of the vesicle yields an accurate representation of an authentic HIV-1 liposome.

The overall lipid composition of mature HIV-1 virions is known from precision lipidomics data, but compositions specific to each leaflet are not. Although the initial liposome model used in this work is based on existing knowledge of plasma membrane asymmetry,^24^ it is a hypothesis for the actual lipid distribution. However, during a long, unrestrained production simulation of the full-scale HIV-1 liposome, the initial asymmetry is maintained despite significant occurrences of lipid flip-flop. The latter indicates that the hypothesized distribution is a good approximation of the asymmetric lipid distribution in the viral envelope.

If this were a flat, asymmetric bilayer, then the lipid asymmetry would be lost over time – without anything to break the up/down symmetry of the leaflets, the lipid distribution will relax to a symmetric distribution. However, the 150 nm vesicle is not up/down symmetric, but is curved, with lipids on the outer leaflet under positive curvature, and lipids on the inner leaflet under negative curvature. The distribution of lipids in the HIV-1 liposome appears to match these curvature preferences. Highly unsaturated lipids (especially with small headgroups like PE) have negative spontaneous curvatures, while mixtures of sphingomyelin and cholesterol have positive spontaneous curvature.^46^ Given that HIV-1 buds from specific locations at the plasma membrane and has a lipid composition that is distinct from that of the host cell, this suggests that HIV-1 selects a composition of its envelope to stabilize the virion, by coupling membrane curvature and asymmetric lipid composition.

Further, our lipidomic representation of the HIV-1 vesicle matches the preference of HIV-1 to enrich its bilayer with Phosphatidylserine, Ceramide, Sphingomyelin and Cholesterol, as reported empirically for various cell lines. ^21,22^ The preferential enrichment of distinct constituents of the lipidome is likely to accommodate morphogenesis, through aforementioned coupling of compositional asymmetry to curvature and to provide domains mimicking detergent-resistant membranes (DRM) with which the Gag polyprotein is known to associate.^21^

Viral entry requires the coordinated action of the envelope protein and CD4+ receptors in order to fuse the host cell membrane and the viral envelope. Furthermore, the spatial distribution of the HIV-1 envelope glycoprotein on the surface of the virion is dependent on stage of the life cycle of the virus. ^47,48^ Whether lipids play a role in organizing the viral envelope for fusion or during maturation is unknown. ^47^ However, the observation of fluctuating nanodomains on the (protein-free) viral liposome indicates that this may indeed be the case. These domains are roughly 10 nm in diameter and enriched in sphingolipids, perhaps providing a dynamic platform for the organization of the envelope protein during maturation and fusion.

Further, experimental evidence suggests that the presence of Cholesterol in biological membranes leads to entropy driven phase separation of L_*d*_ and L_*o*_ domains, ^49,50^ where Cholesterol participates in either phase without preference to serve the end of hydrophobic tail packing in highly ordered membrane environments. These distinct phases are responsible for heterogeneous lateral diffusion along membrane surfaces. The sphingolipid-enriched, Cholesterol-containing domains observed in the present work support this experimental finding. Aside from their potential role in mediating the localization of envelope proteins throughout viral maturation, these domains may serve an additional entropic purpose to aid the packing of lipids within the outer leaflet, where heterogeneous lateral diffusion was observed (Fig. 5). Additionally, tight packing of lipids is known to lead to slow lateral diffusion, ^49^ which may explain the distinct difference in lateral mobility observed between the outer and inner leaflets in the HIV-1 liposome, where packing is tighter in the positively curved outer leaflet than in the inner leaflet. This hetereogenous mobility would have an impact on the lateral diffusion of the envelope protein as it would encounter two different lipid environments, namely the outer leaflet by the receptor binding domain of gp41 and the interior leaftlet by its cytoplasmic tail.

Altogether, our study provides insight into the molecular behavior of HIV liposomes with a high level of detail. The lipid vesicle of HIV contains an asymmetric composition of lipids across monolayers, asymmetry which was persistent throughout molecular dynamics simulation. Together with heterogeneous lateral diffusion and the observed transbilayer diffusion, the persistence of macroscopic qualities of the liposome indicates a central role for the vesicle, and its composition, in the HIV viral replication cycle, where such properties are maintained with the intent of stabilizing the virion through maturation, supporting viral protein components and enabling infectivity.

## Methods

### Vesicle Construction

The approximately 300,000 lipid molecule positions were initially seeded in two spheres, representing leaflets, using the packmol library. ^51^ The relative abundances of the lipid species were based on the average per-virion values obtained in previous lipidomics experiments for HIV-1. ^21,22^ Molecular placement with packmol did not fully converge, so an alchemical phasing procedure was used to progressively expand the effective CG particle radii to relax steric conflicts gradually.^52^ A 1000 frame alchemical growth trajectory spanning *λ* = 0 to *λ* = 1 was visually inspected at every 10th frame to determine the point at which lipids had deviated too far from the HIV-1 vesicle because of steric conflicts during lambda-expansion. The acceptable movements were capped at *λ* = 0.2 (frame 20) and all twenty coordinate sets in that set of frames were energy minimized to assess viability (*F*_*max*_ < 10^4^). The least disruptive (lowest) *λ* value at which coordinates were viable for energy minimization was frame 7 (*λ* = 0.08), and these coordinates were selected for preparation of a production simulation.

The selected lipid vesicle starting configuration was placed in a rhombic dodecahedron container using GROMACS 4.6.5 with a 3.5 nm distance buffer between the vesicle and the simulation container boundary. The dodecahedral system was hydrated with MARTINI water, using a random seed and a van der Waals radius of 0.24 nm. In total, 20541842 water particles were added by GROMACS, and the hydrated system was successfully energy-minimized using the steepest descent algorithm in GROMACS. The simulation system was neutralized by replacing 25,000 MARTINI water molecules with NA+ ions, followed by another successful steepest descent energy minimization. The final solvent preparation step was to convert to 5% antifreeze particle composition, as described previously. ^26^ After confirming that the energy minimized solvated coordinates were sensible based on visual inspection, the system was prepared for long-term (production) simulations with GROMACS 2016.

After approximately 1.5 *μ*s of equilibration, we noticed two issues with the HIV-1 vesicle system: there were a few free-floating lipids inside and outside of the ultrastructure, and double bilayer anomalies were present, i.e., regions with more than two lipid leaflets had formed. The bilayer anomalies were excised using a combination of the DBSCAN algorithm^53^ in scikit-learn^54^ and in-house code to reintegrate false-positive selections.

Following the surgical procedure to remove bilayer anomalies, the vesicular system was rehydrated using GROMACS 4.6.5, with MARTINI water using a van der Waals radius of 0.24 nm. 20,700,856 water particles were added and the hydrated system was energy minimized using the steepest descent algorithm. The hydrated system was neutralized with 22,465 MARTINI sodium ions and again energy minimized with steepest descent. The solvent was converted to 5 % antifreeze particles as described above, leaving 19,644,472 conventional waters in the system, and again successfully energy minimized with steepest descent. Following visual inspection of the vesicular system, the extended production simulation was prepared as before using GROMACS 2016.1.

After an additional 1.5 *μ*s of post-surgery vesicle simulation, there was a gap in computational resource availability, and when additional resources were finally secured, we repeated the simulation preparation procedure a third time, this time rehydrating from the final (dehydrated) 1.5 *μ*s post-surgery equilibration snapshot that was still available. 21,666,258 MARTINI waters and 22,465 NA+ ions were added, and then solvent was converted to 5% antifreeze particles (1,082,189 added in place of water). Intervening energy minimizations were as described above. To maximize performance, we prepared our long-term production simulation using GROMACS 2018.1.

### Simulation Parameters

In addition to the lipid components, the system was solvated with 5% antifreeze particles (1,082,189 MARTINI WF residues), 95% water particles (20,561,604 MARTINI W residues), and 22,465 Na ions, for a total of 24,585,206 beads.

**Table 1:** Summary of HIV-1 vesicle lipid composition. Lipid names match those used in the MARTINI 2.1 forcefield. There are 24 lipid species in total.

| Lipid Name | Number of Molecules |
| --- | --- |
| POPS | 1161 |
| DLPC | 4929 |
| PRPS | 8931 |
| DPCE | 159 |
| PNSM | 10542 |
| POPC | 10128 |
| POPE | 1092 |
| DOPC | 2855 |
| PQPE | 583 |
| PGPS | 5579 |
| PQPS | 1584 |
| DPPC | 6754 |
| PIPE | 9084 |
| PRPE | 9823 |
| CHOL | 129019 |
| PAPE | 13182 |
| PIPS | 3651 |
| DPSM | 43334 |
| PAPS | 1559 |
| PUPE | 15531 |
| DPGS | 592 |
| PUPC | 776 |
| PRPC | 518 |
| PGPE | 1833 |

All simulations were performed using the MARTINI 2.1 coarse-grain forcefieldi. ^30^ After the 1.5 *μ*s initial equilibration using GROMACS^55^ version 2016.4, 5.2 *μ*s production simulations were performed using GROMACS 2018.1 with 10 fs timesteps, the Verlet cutoff scheme, and reaction field electrostatics. Isotropic pressure coupling was employed using the Berendsen barostat at a time constant of 20 ps, compressibility of 1 × 10^−6^ bar^−1^, and reference pressure of 1.0 bar. Lipids and solvent were separately temperature coupled with a 323 K reference temperature using the Berendsen thermostat, and a 1 ps time constant.

### Analysis of Simulations

Trajectory data files were read and exposed to the Python interpreter using the MDAnalysis library. ^56,57^ Trajectories were visualized using VMD,^58^ and analysis plots were produced using matplotlib. ^59^ Array-based calculations of simulation properties leveraged the NumPy library. ^60^ The SciPy library^61^ was used for a number of scientific algorithms.

#### Sphericity Tracking

Sphericity of the liposome was calculated using the ratio between surface area and volume previously reported in a geological context.^62^ The surface area of the vesicle was estimated by recasting the Cartesian coordinates of the headgroups to spherical coordinates and taking the average of the radius of the system centered at the origin. Based on the sphericity tracking analysis during equilibration, the surface area was estimated for the circumference encompassing the average radius of the headgroups of both monolayers.

### Local bilayer thickness

Local bilayer thickness was computed as the distance between centroids of headgroups in regions of radius 50 Å on each leaflet surface. To ensure uniformity of sampled regions on the vesicle surface, a Poisson disk sampler developed by the authors^32^ was employed.

### Lipid translational and trans-bilayer diffusion

The translational diffusion coefficients of the lipids in each leaflet were calculated using the approach previously reported for viral simulations ^27^ using,

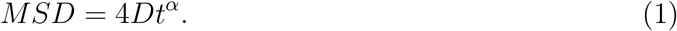

Where MSD represents the mean square displacement (MSD) of the centroid of every lipid, *D* is the lateral diffusion coefficient and *α* is an arbitrary scaling factor. *D* and *α* were estimated using linear least-square fitting of MSD for simulation windows of size: 5, 25, 50, 100, 250, 500, 1000 ns. The standard deviations for the diffusion coefficient and the scaling factor were derived from the square root of the covariance matrix of the linear least-square fitting as previously described. ^27^

Analysis of trans-bilayer diffusion measured the number of flip-flop events per unit of time, for each lipid species, using a 3D space classifier developed by the authors. ^32^ Using lipid tail groups, the volume of the vesicle system was classified such that molecular density corresponding to the tail group selection served as a barrier separating interior and exterior regions of the system, representing the inner and outer leaflets, respectively. For each frame of the trajectory, headgroups were characterized according to their positions within the classified volume, i.e., interior or exterior. The latter was made possible through a massively parallel analysis framework. ^29^ To mitigate error resulting from undulations of the vesicle, volume classification was repeated every 20 ns throughout the time series analysis. Data resulting from headgroup tracking were used to quantify inner-to-outer and outer-to-inner translocation events and determine effective rates thereof for each species.

### Mobility analysis

Lateral mobility of lipids was measured by estimating the root mean square fluctuation (RMSF) as 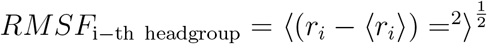; where 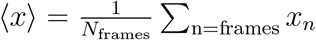. RMSF values were calculated for lipid headgroups over a simulation window of size of 1 *μ*s. Results are reported as the mean RMSF for headgroups of each lipid in the vesicle.

### Bending rigidity

The bending rigidity of the full-scale vesicle was calculated from a well-established spherical harmonic analysis for spherical liposomes. ^38^ First, a single reference frame is defined by estimating the undulating radial surface (URS).^38^ The initial URS is estimated by recasting a *ψ,θ*-grid from the coordinate system of the vesicle, where *θ* ∈ [0*, ψ*] and *ψ* ∈ [0, 2*π*], for the colatitude and longitude, respectively. A total of two inital grids were generated, each one corresponding to one monolayer of the vesicle. The angular resolution for the two grids was *dθ* = *dψ* = 0.042 rad. based on a cutoff wavenumber of 0.5 nm^−1^ and a cutoff filter of 2.5 nm^−1^ to mitigate discontinuities at the poles of the vesicle. ^38^ After defining the surface of the inner (*r*_*in*_(*ϕ, ψ*)) and outer monolayer (*r*_*out*_(*ϕ, ψ*)), the undulating surface *r*_*und*_ was calculated for the vesicle as shown in equation 2.

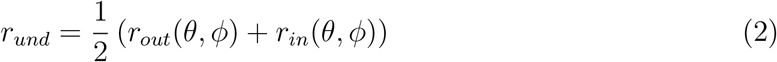

The average radius of the undulating surface 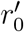 was then used to define the normalized radial fluctuation as

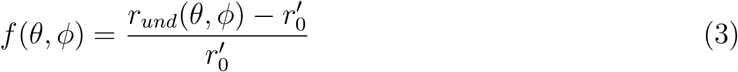

Subsequently, spherical harmonic analysis was employed to decompose the normalized radial fluctuations from equation 3. Liposome fluctuations were expanded in spherical harmonics of degree *l* and order *m* as

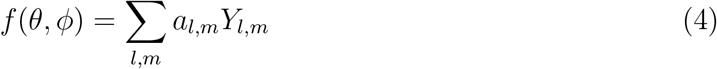

The linear combination of spherical harmonics from equation 4 consists of harmonic coefficients *a*_*l,m*_ and basis functions 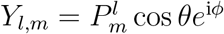; where 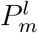 are Legendre polynomials. The order and degree of *Y*_*l,m*_ are dependent on the number of points describing the initial undulating surface. In the case of the HIV-1 vesicle, the spherical harmonics were expanded as a linear combination of degree 74 and order 149.

Finally, the pseudo-inverse of the matrix *Y* composed by the basis functions is used to estimate the harmonic coefficients *a*_*l,m*_. The *a*_*lm*_ can be used to obtain the power spectra of undulations by fitting the harmonic coefficients to the Helfrich continuum model for undulations on a sphere with vanishing spontaneous curvature,^63^ shown in equation 5. The fitting yields an estimation of the bending rigidity *k*_*c*_ at temperature *T* as

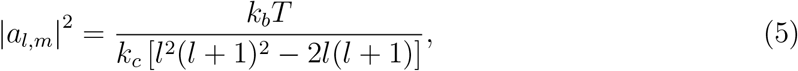

where *k*_*b*_ is the Boltzmann constant.

## Acknowledgments

The authors acknowledge funding from the US National Institutes of Health award P50AI1504817 and P20GM104316. Methodology development for the present work was provided by the National Science Foundation award MCB-2027096, funded in part by Delaware Established Program to Stimulate Competitive Research (EPSCoR). This research is part of the Frontera computing project at the Texas Advanced Computing Center. Frontera is made possible by NSF award OAC-1818253. This work used the Extreme Science and Engineering Discovery Environment, which is supported by the National Science Foundation (Grant ACI-1548562). This work used XSEDE Bridges and Stampede2 at the Pittsburgh Super Computing Center and Texas Advanced Computing Center, respectively, through allocation MCB170096. Computational resources were also provided by a LANL institutional computing grant on the Grizzly Supercomputer (2019-2020), and a Great Lakes Consortium for Petascale Computation grant for the NCSA Blue Waters supercomputer (2018-2019). The authors thank Fabio Gonzalez for insightful conversations and technical support.

## Supporting Information Available

**Figure S1:**
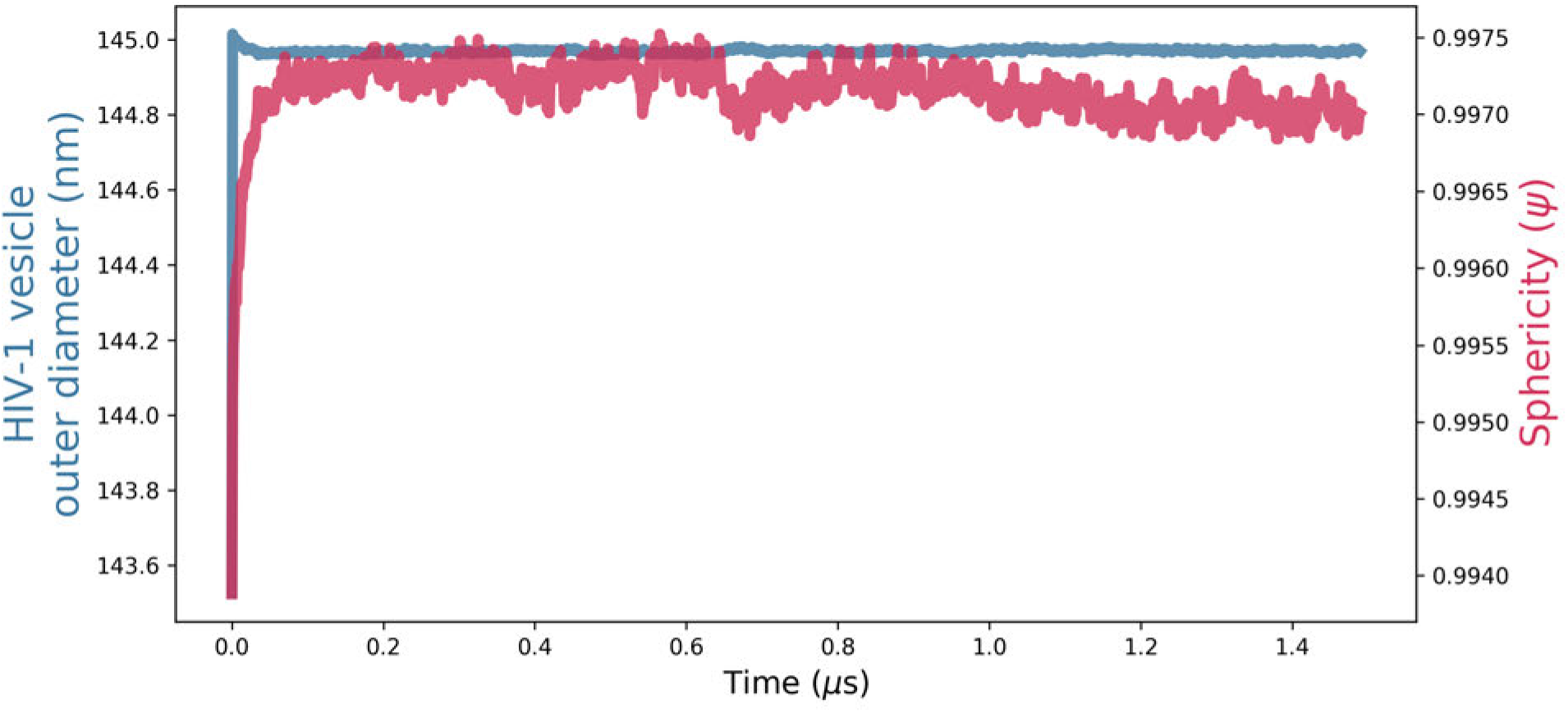
Tracking outer diameter and sphericity of the HIV-1 vesicle simulation over the course of an initial 1.5 *μ*s equilibration, another 1.5 *μ*s post-surgery equilibration, and an extended production simulation.

**Table S1:** Summary of HIV-1 vesicle inner leaflet and outer leaflet lipidome. Values represent the number of molecules in the inner and outer layer.

|  | Time: 0 ns |  | Time: 5200 ns |  |
| --- | --- | --- | --- | --- |
| **Lipidome** | **Inner leaflet** | **Outer leaflet** | **Inner leaflet** | **Outer leaflet** |
| **CHOL** | 43390 | 83481 | 42612 | 84618 |
| **DPCE** | 0 | 159 | 2 | 157 |
| **DPSM** | 194 | 43140 | 194 | 43142 |
| **PGPE** | 1833 | 4 | 1832 | 6 |
| **PIPS** | 3640 | 23 | 3640 | 22 |
| **POPE** | 1091 | 2 | 1090 | 6 |
| **PQPS** | 1581 | 9 | 1581 | 9 |
| **PRPS** | 8912 | 54 | 8909 | 72 |
| **DLPC** | 25 | 4904 | 25 | 4904 |
| **DPGS** | 7 | 585 | 7 | 585 |
| **PAPE** | 13163 | 59 | 13159 | 77 |
| **PGPS** | 5572 | 17 | 5569 | 20 |
| **PNSM** | 42 | 10500 | 42 | 10500 |
| **POPS** | 1160 | 4 | 1160 | 4 |
| **PRPC** | 2 | 516 | 2 | 516 |
| **PUPC** | 1 | 775 | 1 | 775 |
| **DOPC** | 13 | 2842 | 13 | 2842 |
| **DPPC** | 29 | 6725 | 29 | 6725 |
| **PAPS** | 1557 | 7 | 1555 | 6 |
| **PIPE** | 9081 | 34 | 9077 | 36 |
| **POPC** | 34 | 10094 | 34 | 10094 |
| **PQPE** | 582 | 4 | 582 | 4 |
| **PRPE** | 9808 | 61 | 9805 | 59 |
| **PUPE** | 15516 | 64 | 15507 | 101 |

**Table S2:**
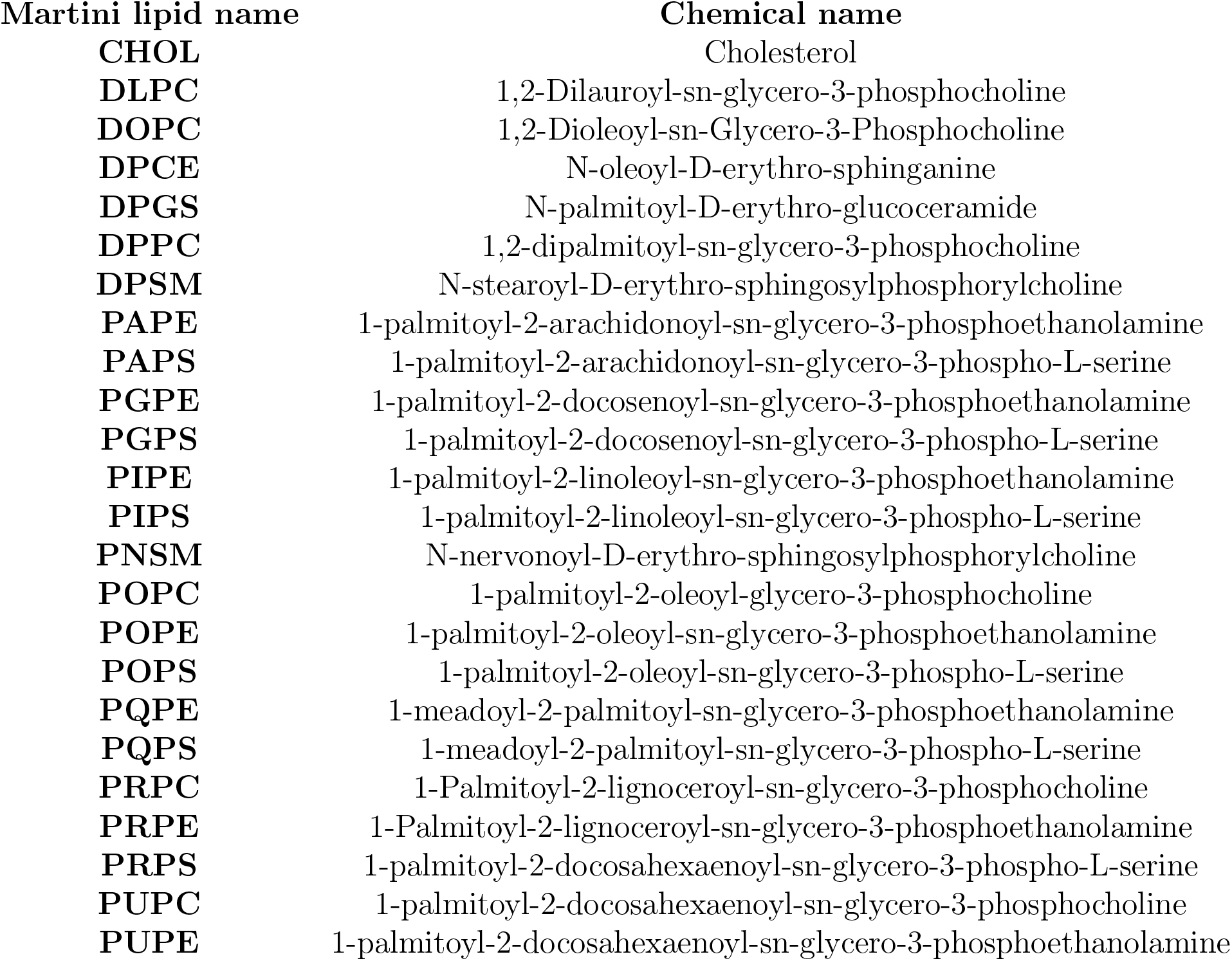
Chemical names and MARTINI residue names of the lipids used for the vesicle model

**Figure S2:**
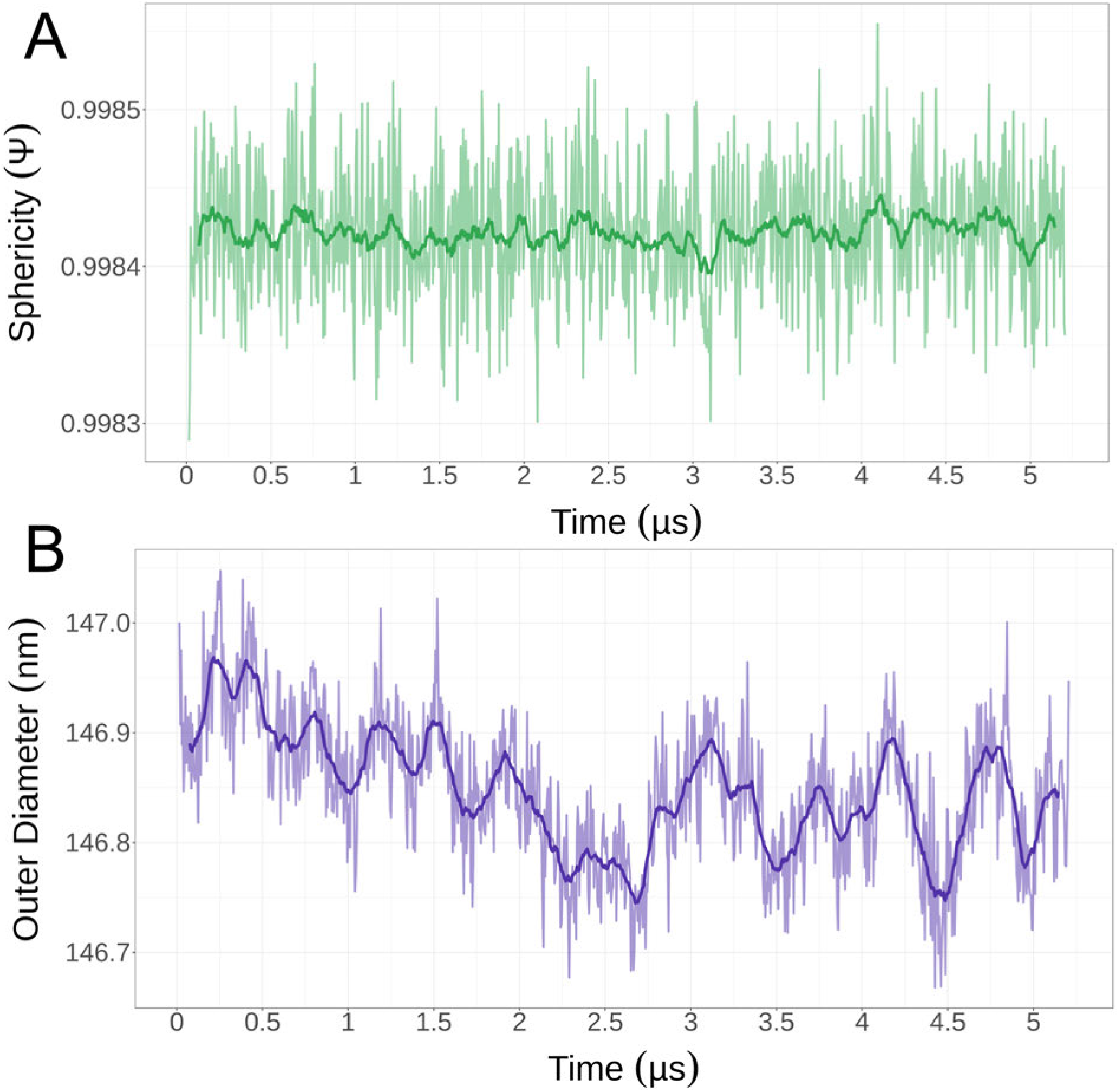
Tracking sphericity (**A**) and outer diameter (**B**) of the HIV-1 vesicle over the course of 5.2 *μ*s production simulation. The darkened line traces represent a rolling mean, computed with a window size of 0.125 *μ*s for both analyses.

**Figure S3:**
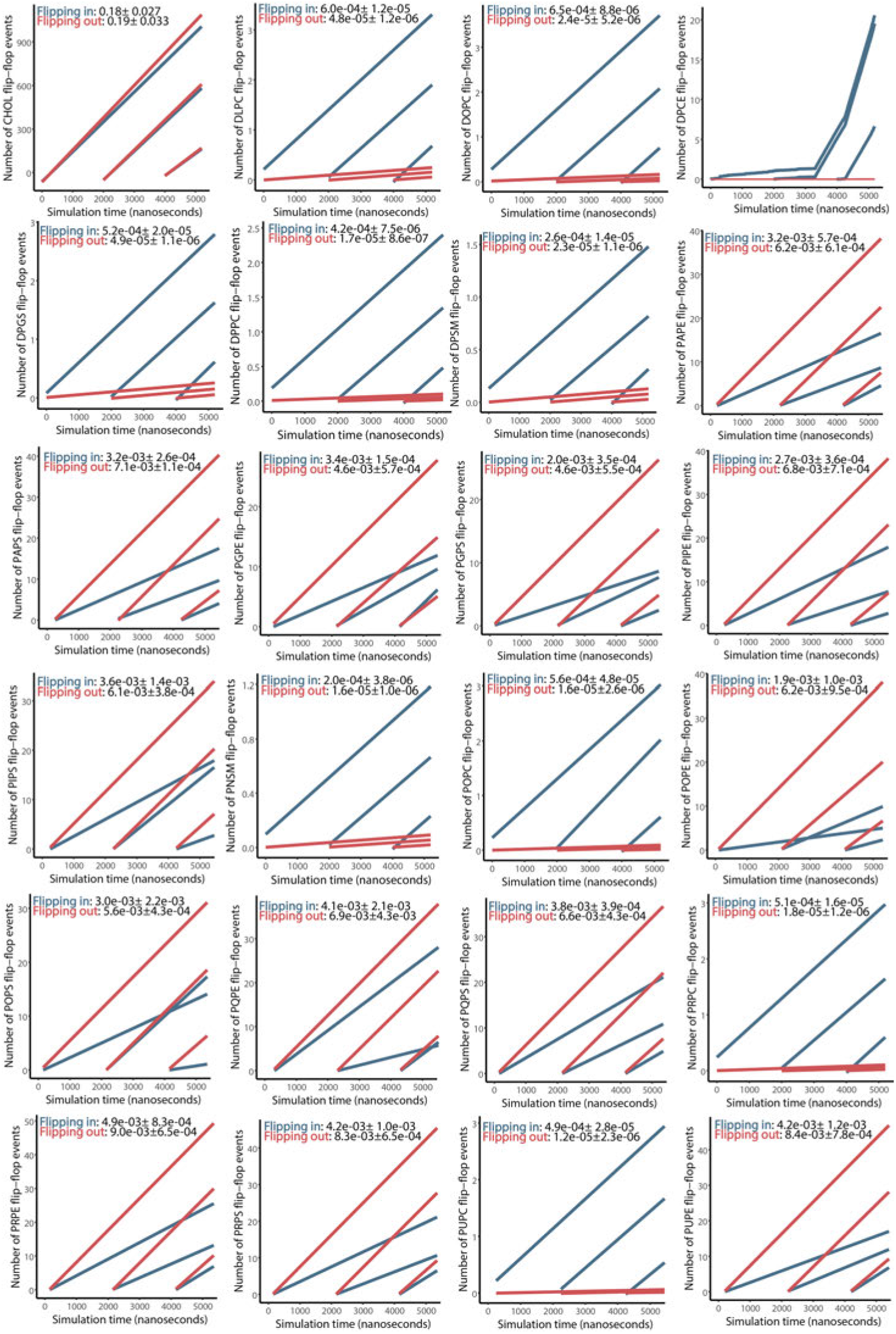
Lipid flip-in and flip-out rates during the 5.2 *μ*s of MD production of the HIV-1 vesicle. The rates were estimated for translocation to the inner and outer leaflet, shown in red and blue, respectively. The number of events was normalized against the number of each lipid type. The rates were estimated as a linear fit to the number of events at three timescales, for both trans-bilayer diffusion events. The rates of DPCE were not computed due to the linear fit obtained.

**Figure S4:**
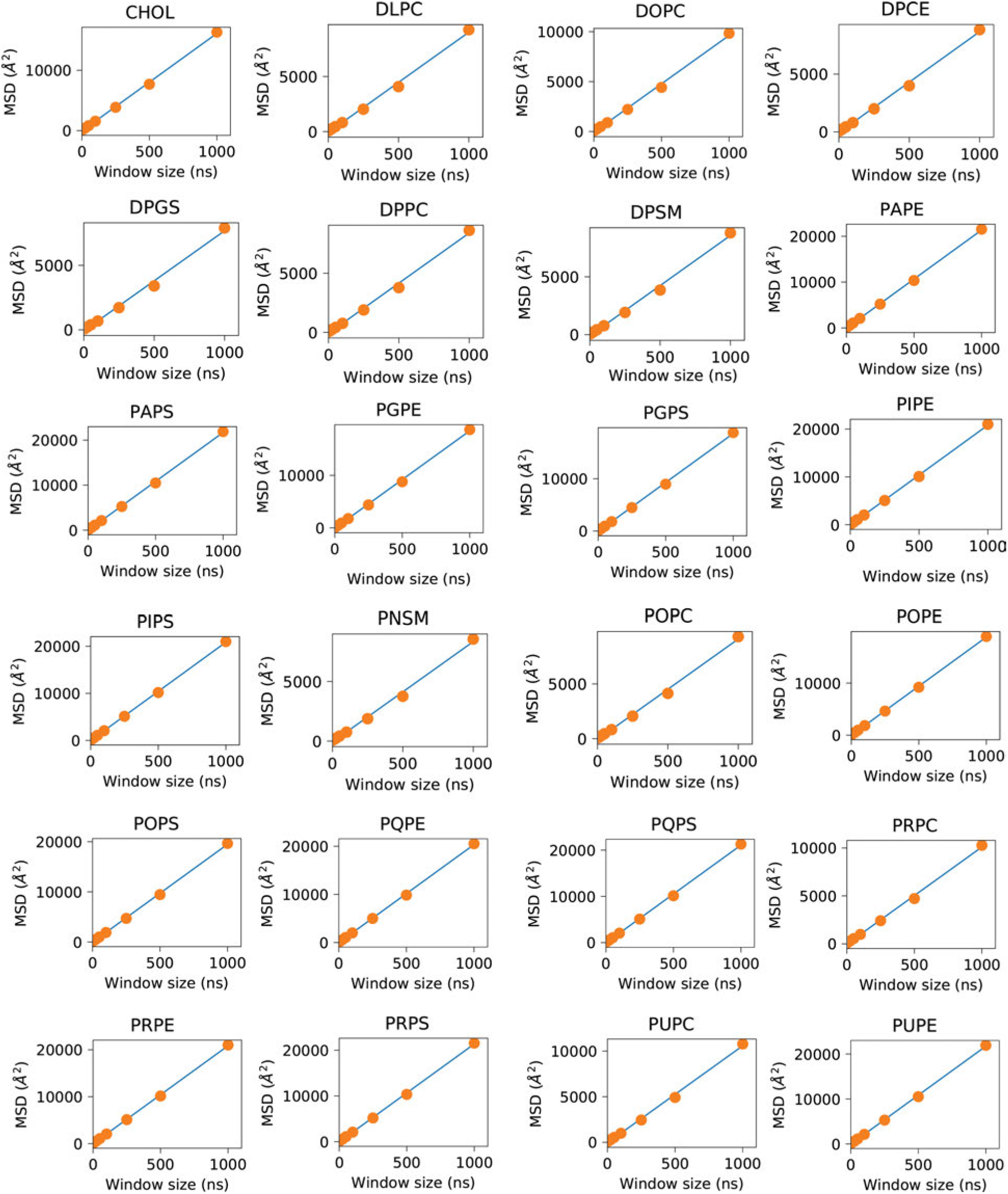
Linear fit of the mean square displacement (MSD) analysis at window sizes 0, 50, 100, 250, 500, 1000ns for each lipid type, from 5.2 *μ*s simulation of the vesicle.

**Figure S5:**
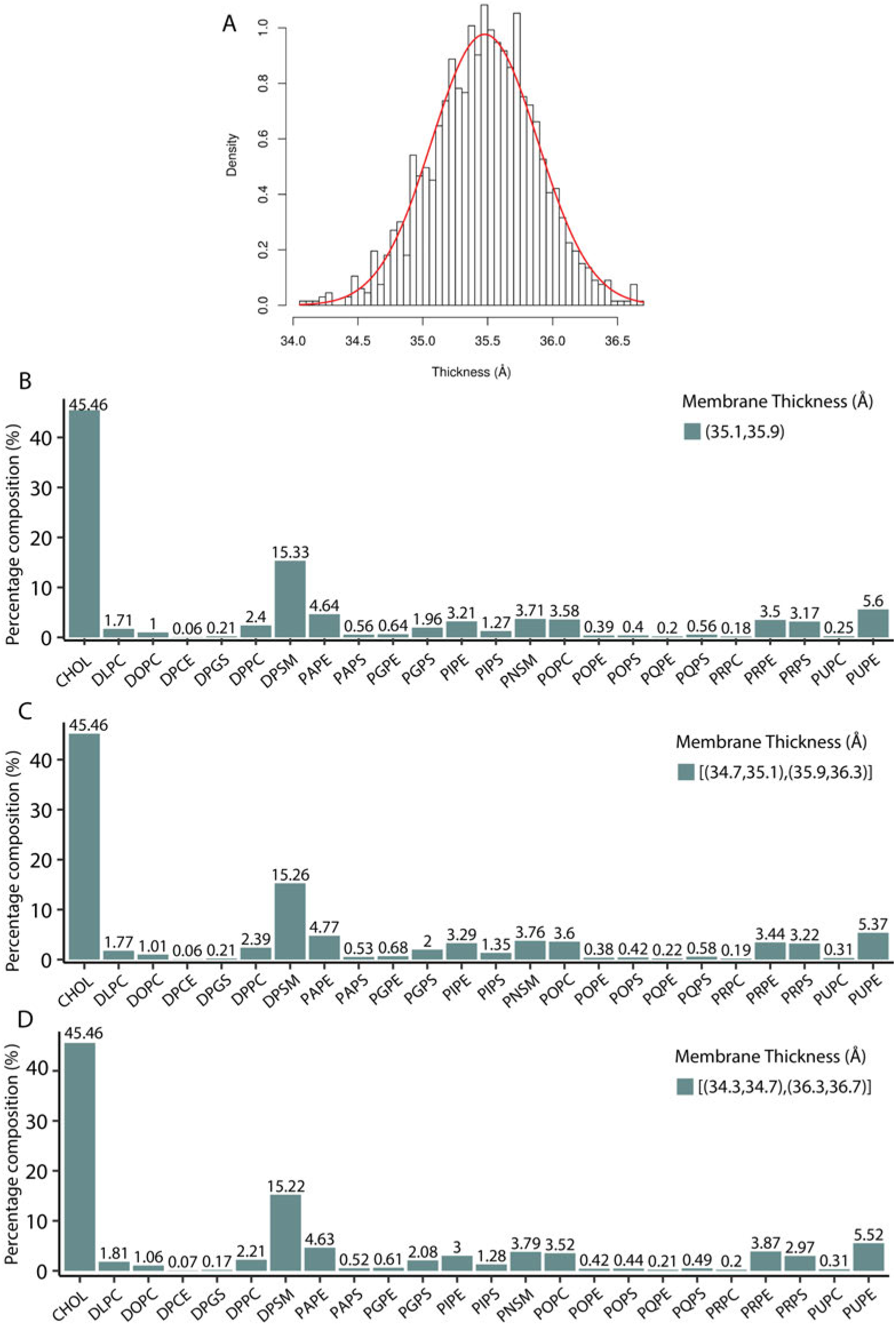
Percent composition of the vesicle at three different intervals, according to the Gaussian distribution of bilayer thickness values

**Figure S6:**
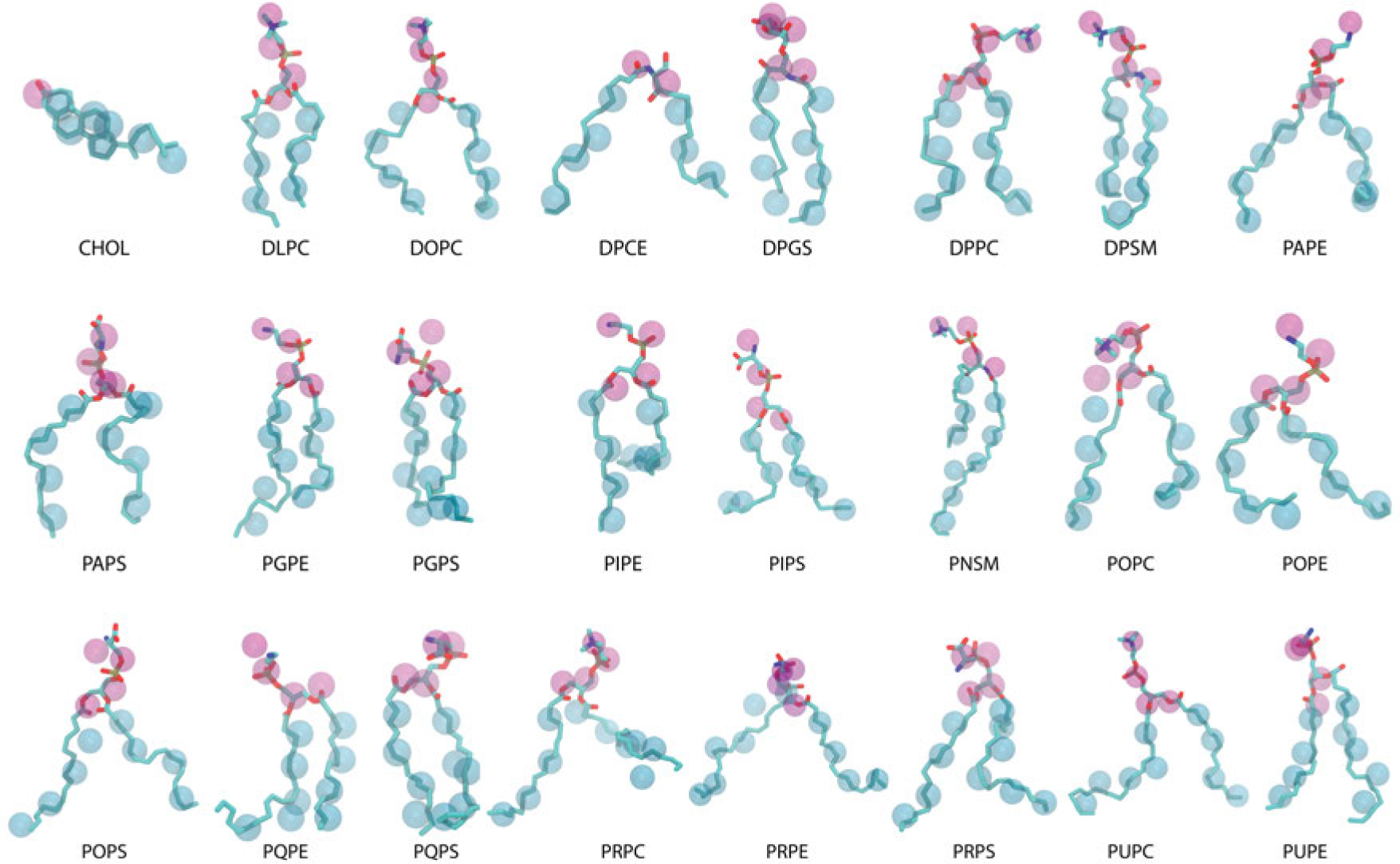
Lipidome of the vesicle used in the present study. Lipids are represented as a superposition of the CG beads and the atomistic model. Transparent CG beads display the headgroups in magenta and tails in cyan.

**Figure S7:**
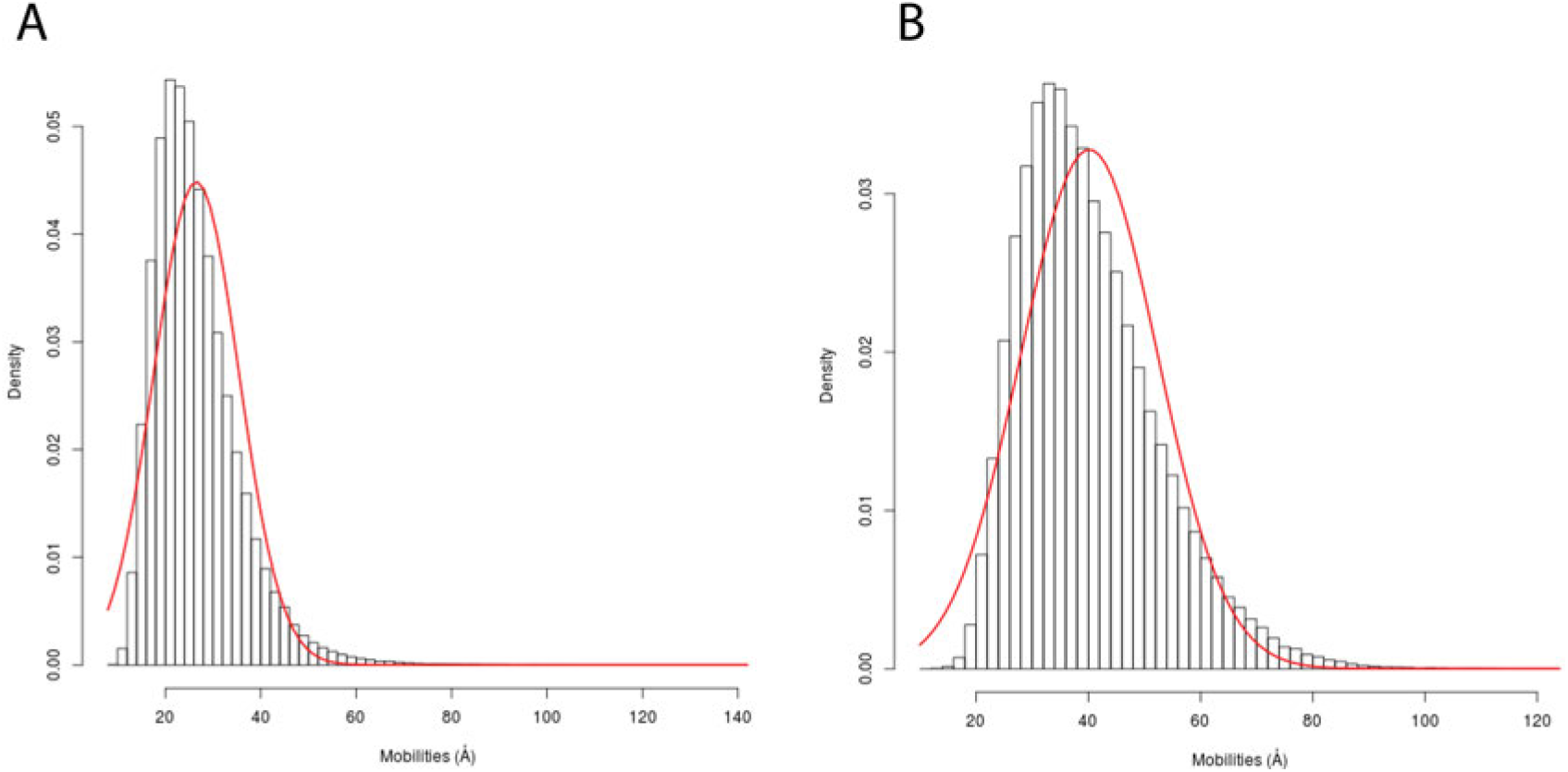
Distribution of lipid headgroup mobility and Gaussian fit (red line) for 500 ns of MD production. **A** Mobility distribution of lipids in the inner monolayer. **B** Mobility distribution of lipids in the outer monolayer.

**Movie M1:**
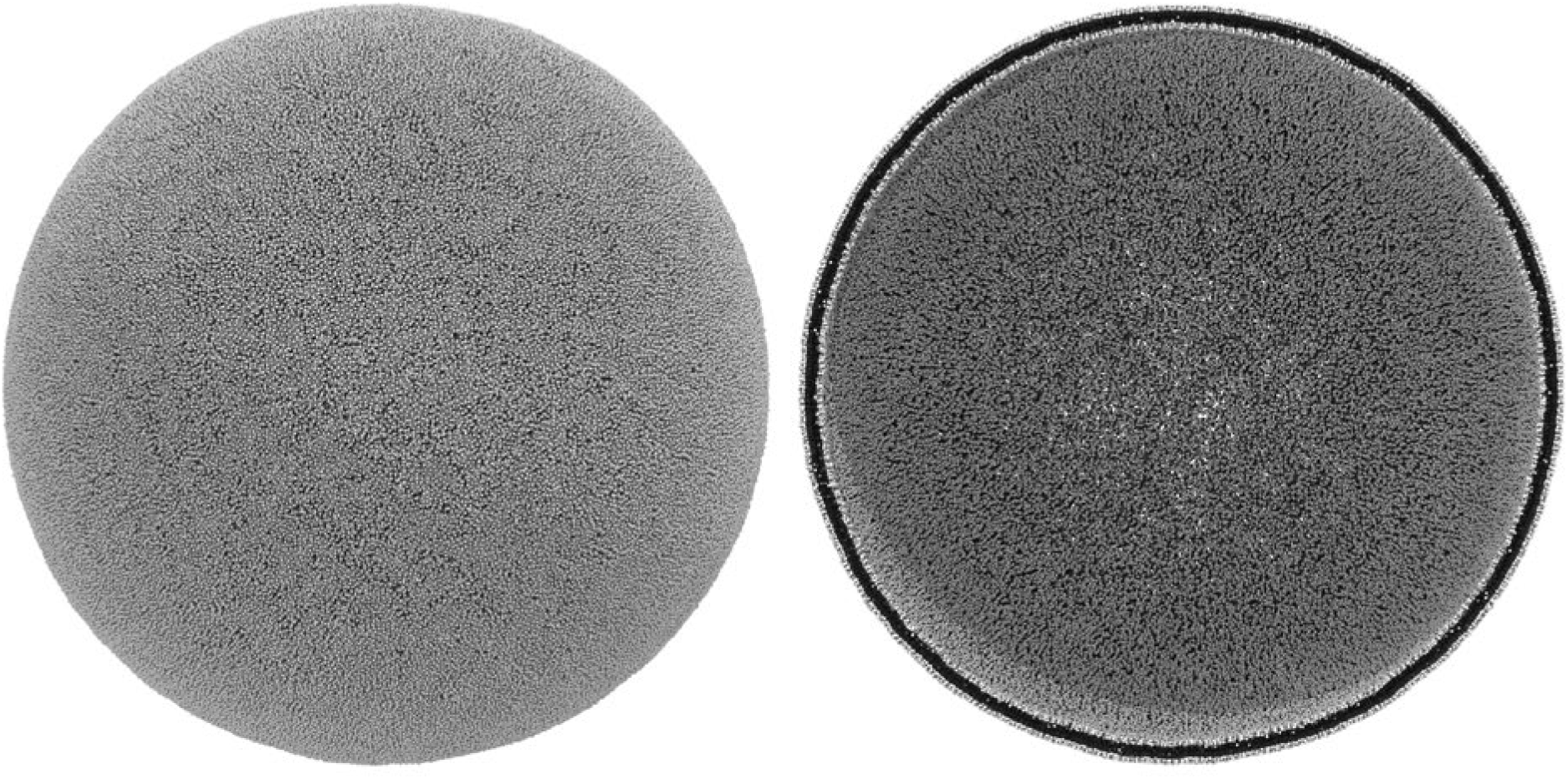
Molecular dynamics simulation of a full-scale HIV-1 lipid vesicle.

**Movie M2:**
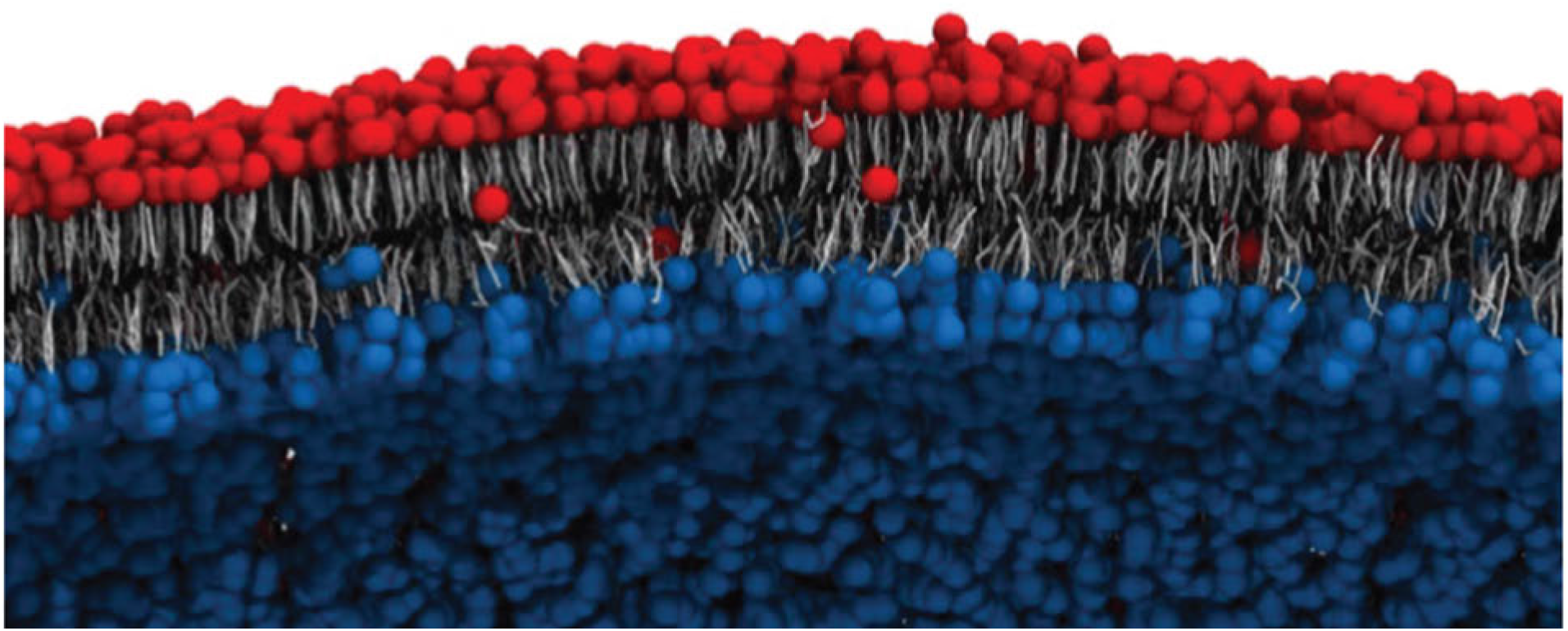
Visualization of the flip-flop of lipids between the inner (blue) and outer (red) leaflet.

**Movie M3:**
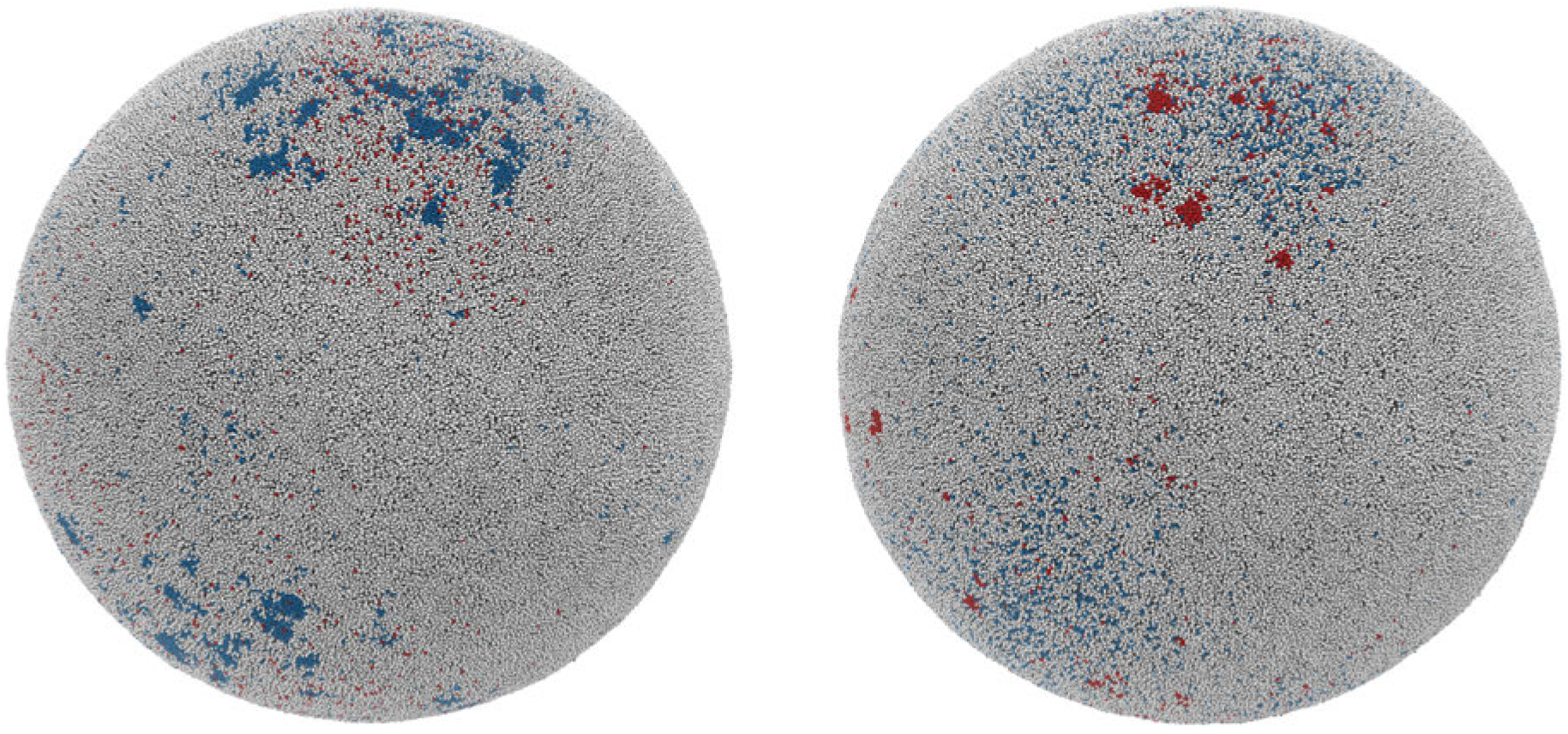
Identification of high and low mobility regions. The initial low mobility regions (blue) are transient, as are the formation of low mobility regions (red) evolving throughout production MD simulation. Overall the simulations indicate that the formation and dissipation of high and low mobility regions is dynamic.

## Notes

### Competing Interest Statement

The authors have declared no competing interest.

